# Homologous recombination and transposon propagation shapes the population structure of an organism from the deep subsurface with minimal metabolism

**DOI:** 10.1101/249516

**Authors:** Alexander J Probst, Jillian F. Banfield

## Abstract

DPANN archaea are primarily known based on genomes from metagenomes and single cells. We reconstructed a complete population genome for *Candidatus* “Forterrea”, a Diapherotrite with a predicted symbiotic lifestyle probably centered around nucleotide metabolism and RuBisCO. Genome-wide analysis of sequence variation provided insights into the processes that shape its population structure in the deep subsurface. The genome contains many transposons, yet reconstruction of a complete genome from a short-read insert dataset was possible because most occurred only in some individuals. Accuracy of the final reconstruction could be verified because the genome displays the pattern of cumulative GC skew known for some archaea but more typically associated with bacteria. Sequence variation is highly localized, and most pronounced around transposons and relatively close to the origin of replication. Patterns of variation are best explained by homologous recombination, a process previously not described for DPANN archaea.

---

The DPANN archaea comprise the Diapherotrites, Parvarchaeota, Aenigmarchaeota, Nanoarchaeota and Nanohaloarchaeota (Rinke et al. 2013) and other archaea (Castelle et al. 2015). Many have rRNA genes sequences that are not amplified by commonly used primers, so they are often missed in gene surveys (Eloe-Fadrosh et al. 2016). DPANN are members of microbial communities across a diversity of Earth’s environments. All have small, enigmatic genomes with many hypothetical proteins. Most are inferred to adopt symbiotic lifestyles based on missing metabolic capacities or experimental analyses. However, all members of the Diapherotrites phylum have been suggested to be free living (Youssef et al. 2015; Castelle et al. 2015).

CO_2_-driven eruptions of Crystal Geyser (Utah, USA) deliver to the surface groundwater from three different depth (aquifers hosted in the Navajo, Wingate and the Entrada sandstone formations) (Probst et al. 2018). In water samples deriving from intermediate and deeper aquifer regions (**Fig. S1**) we identified a novel archaeon for which we recovered a near-complete draft genome. The genomic dataset from sample CG_2015-17, initially comprising five scaffolds, was selected for improvement. The draft genome was subjected to seven rounds of curation that involved mapping of reads from the CG_2015-17 dataset to the scaffold set using bowtie2 (Langmead & Salzberg 2012) (‐‐sensitive). Scaffold ends were extended based on the newly recruited reads and their unplaced paired reads using Geneious software. Mapped reads were also used to close internal scaffold gaps and generate contiguous sequences (contigs) and to remove local assembly errors. Three additional small contigs (1.3 – 3.4 kbp) were identified based on identity with extended contig ends and added to the dataset and a small stretch of candidate sequence was identified from sample CG_2015-09. Inclusion of the added sequences was verified in the subsequent rounds of read mapping-based curation. One 1.5 kbp contig was identified as a copy of a transposase and thus excluded. Ultimately, five contiguous sequences were combined into one sequence in the only way that enabled inclusion of all sequences into a circular genome (**Fig. 1A and S2**). The final genome has one exactly duplicated transpose. Assembly of reads mapped to the transposon sequence flanks re-generated all paths containing transposons (both the two that occurred in all cells as well as all points of insertion in a subset of cells). Thus, we verified that no additional genome fragments flanked by transposon sequences were unaccounted for. Following closure of the genome, we tested for and identified the typical origin to terminus pattern of cumulative GC skew (**Fig. 1B**) that is found in most bacteria and some archaea (Myllykallio et al. 2000). Aberrations in this pattern could indicate a genome assembly error. When the origin was adjusted to the position predicted by the skew pattern, it fell adjacent to a cdc6 gene, as expected for archaea (Myllykallio et al. 2000). The origin position was confirmed using the Z-curve method (Zhang & Zhang 2005) (**Fig. S3**).

**Fig. 1.**
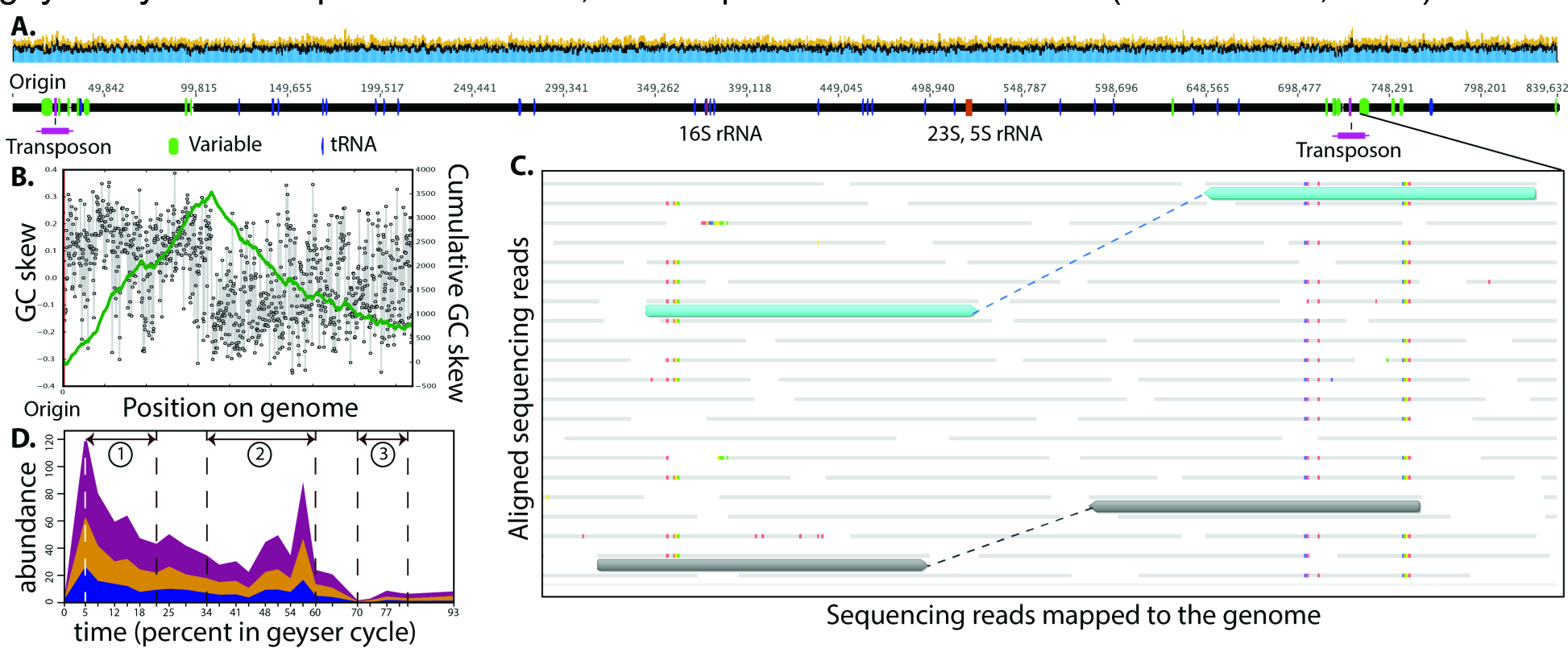
**A.** Overview of the reconstructed, complete genome sequence showing consistent genome coverage by mapped reads (top panel), locations of tRNA and rRNA genes, transposon positions and hot spots of sequence variation (SNPs and indels flanking one tranposon). For details, see the mapped reads file (http://ggkbase.berkeley.edu/Forterrea/organisms) and assembly overview (**Fig. S2**). **B.** Plot of the cumulative GC skew across the genome. The form of the plot is consistent with accurate assembly, and indicates the origin of replication. **C.** Detailed view of local sequence variation within a putative chitobiase gene (colored bars are SNPs). Grey bars and blue bars underline to two sets of paired reads with distinct patterns of SNP linkage. D. Stackplot of relative abundances of Forterrea and a breakeown of the abundances of the two individual strains throughout the eruption cycle of the Geyser. Numbers indicated different phases of the geyser eruption in which intermediate (***1***), deep (***2***), and shallow (***3***) groundwater is sampled as elucidated in Probst et al., 2018. One percent of the geyser cycle corresponds to 70 min, 25 samples were taken in total (Probst et al., 2018).

We analyzed the mapped read dataset to evaluate forms of strain variation other than transposons and identified regions up to ~6.8 kbps in length with elevated frequencies of single nucleotide polymorphisms (SNPs). Notably, SNPs often comprised two distinct strain patterns that transition smoothly into essentially SNP free regions. In a few cases linkage patterns directly indicate recombination (Tyson et al. 2004) (**Fig. 1C**). We reconstructed both of the strain sequences from a single genomic region and then mapped reads from the 25 samples collected over the eruption cycle to these sequences to track the strain relative abundances. Results show a fairly consistent median abundance ratio of ~1:1.5 when water from the deeper aquifers was sampled (**Fig. 1D**). Most sequence variation is concentrated relatively close to the origin of replication and near to the duplicated transposon sequences that are found in all sampled individuals (**Fig. 1A**). The localized SNP patterns, as well as their concentration relative to the origin and transposases, suggest that diversification of the population arises in large part from homologous recombination.

We investigated differences in DPANN genomes reconstructed from samples collected over more than two days of the eruption cycle to test for population shifts that accompanied changes in groundwater source regions To do this, we compared the curated complete genome from CG_2015-17 with de novo reconstructed genome fragments from samples CG_2015-05, ‐07, - 09, ‐10, ‐14, ‐15, ‐16. Due to the available contiguous sequence lengths, around 465 kbp of contiguous sequence could be compared for all samples, 688 kbp for four samples and 694 kbp for three samples. The sequence over the vast majority of all alignments was 100% identical. The pattern of large identical regions in all genomes and local elevated SNP densities reinforces the deduction that the genomes of these deep subsurface DPANN are shaped by the combination of homologous recombination and selective sweeps that remove some sequence variants.

In each case where a SNP distinguished the de novo assembled sequences we found that the same nucleotide was present as a minor variant in reads from CG-2015-17, with just one exception. We also identified a 279 bp insertion within an 802 bp deletion in sequences from two samples (**Fig 2A**). This variant also occurred in a subset of the cells of the CG-2015-17 population. Interestingly, the variant sequences differ substantially immediately prior to this insertion/deletion, with SNPs concentrated in a predicted intergenic region (**Fig 2A**).

**Fig. 2.**
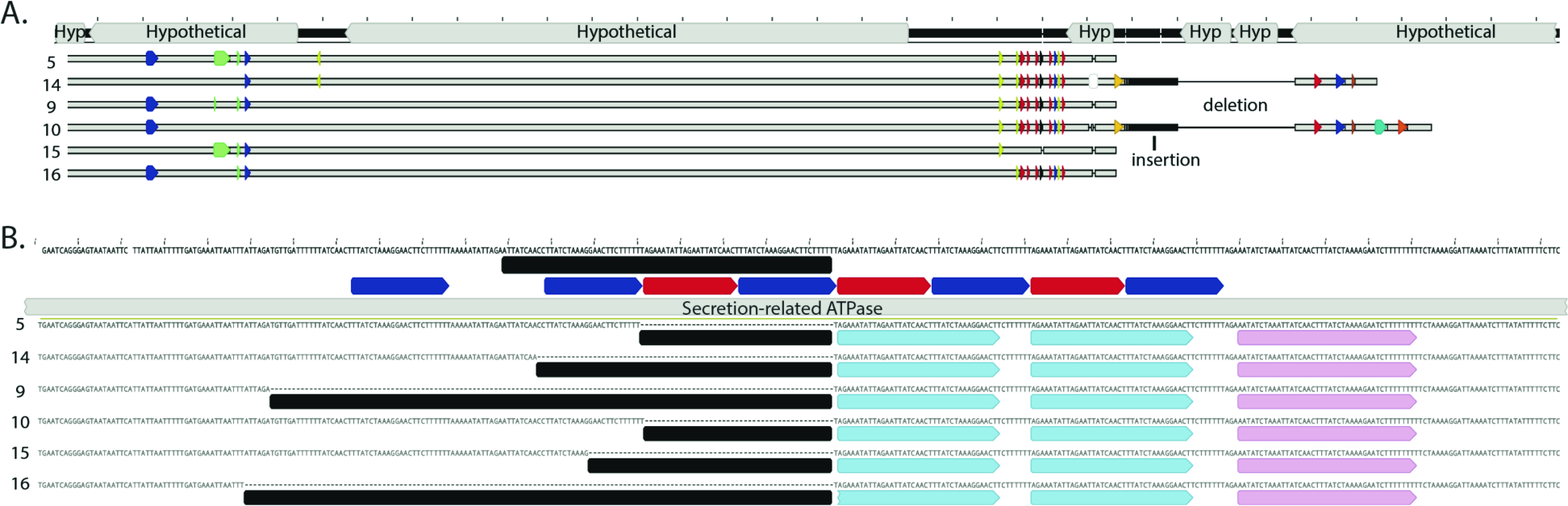
**A.** Comparison of a segment of the curated closed genome sequence from sample 17 with the de novo assembled sequences from other samples. Colored arrows indicate single or a few clustered single nucleotide polymorphisms. Across the entire aligned region of up to ~ 700 kbp (not shown), essentially all polymorphisms and the insertion/deletion events found across samples exist within the population in sample 17. **B.** Diagram comparing the initial and curated sequences of one region within a secretion-related ATPase. Red and blue bars indicate repeated sequences. Black bars indicate blocks of sequence that were omitted in the *de novo* assemblies. Although initial inspection suggested this to be a strain variable region, removal of local scaffolding errors revealed that the sequences in each genome were identical. This result underlines the importance of genome curation prior to strain variation analyses.

A block of inserted or deleted sequence also distinguished sequences from different samples. As this can arise due to local mis-assemblies, we curated all assemblies and showed that all de novo assembled sequences were incorrect, and in a variety of ways. Following manual curation, all sequences agreed perfectly (**Fig. 2B**). This result underlines the importance of curation prior to comparative genomic analyses. As essentially all differences among the genomes sampled over time involve diversity that is evident at a single time point, we attribute cross-sample differences to small shifts in variant abundances.

Over the period studied in the abovementioned comparative analyses, water is continuously drawn from the intermediate aquifer. Thus, the results provide information about population heterogeneity in the aquifer source region. Surprisingly, perhaps, the results point to spatial homogeneity, with source regions dominated by just two main strain types and their recombination-based variants.

The availability of a complete genome enabled analysis of the encoded metabolic potential without concern that missing genes are in gaps. As is typical for genomes of DPANN archaea (Castelle et al. 2015), the predicted proteome is highly novel, with ~45% of all proteins lacking any functional annotation. 19% of the proteins are most similar to those in bacteria (7% are most closely related to proteins of candidate phyla radiation bacteria). The majority of annotated genes function in the information system, encode extracellular proteins or are involved in cell division. Many core metabolic functions, including all components of the electron transport chain, TCA cycle, upper glycolytic pathway, most of the pentose phosphate pathway, synthesis pathways of nucleotides, vitamins and even lipids are absent (**Fig. 3A**). Thus, we infer that the studied archaeon derives most cellular components from a host (**Fig. 3A**).

**Fig. 3.**
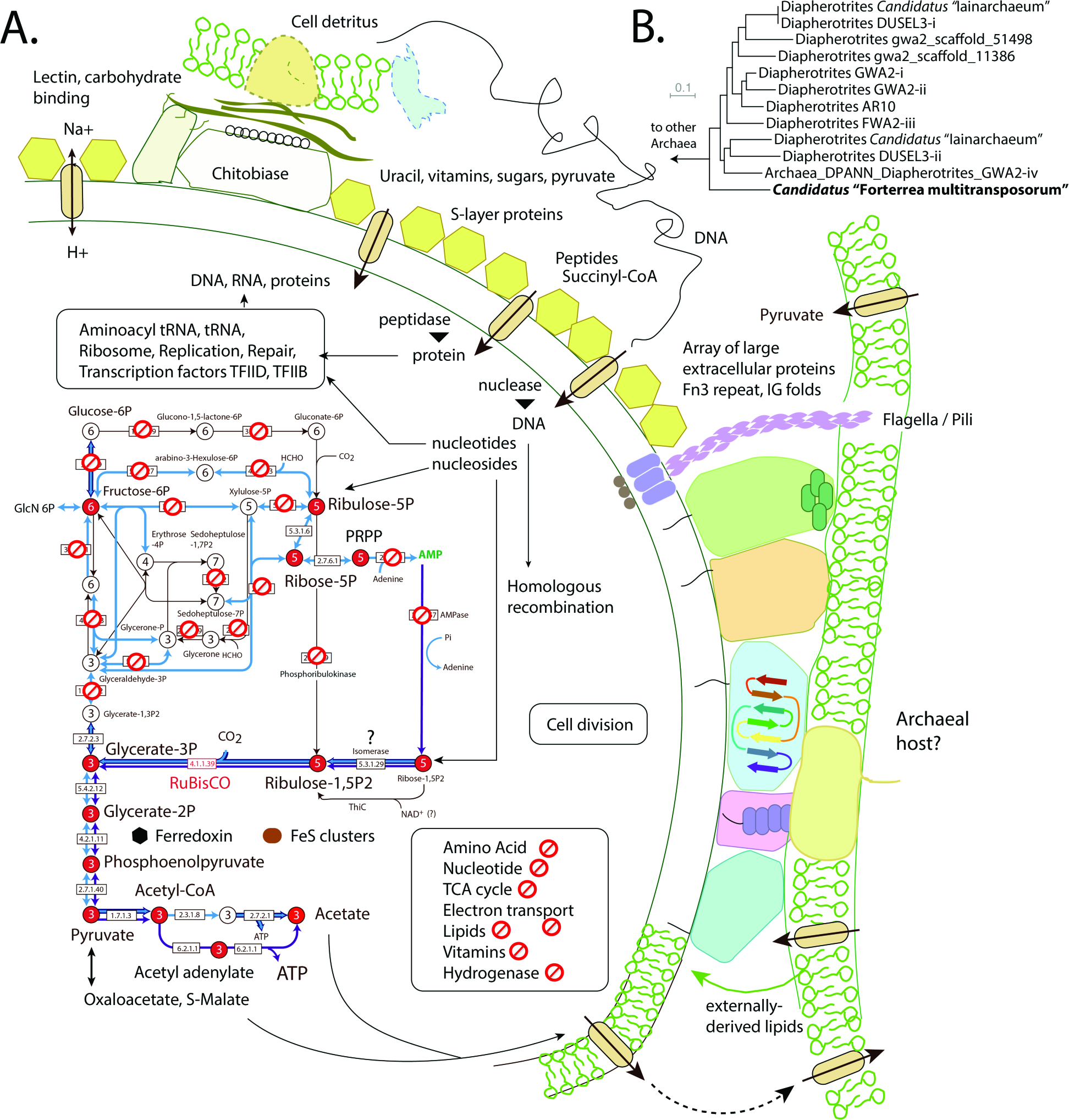
**A.** Cell diagram illustrating metabolic features identified and absent capacities. Large extracellular proteins are predicted to play a role in interaction with other organisms from which many key compounds must be sourced. **B.** 16S rRNA tree placing the archaeon on a deep branch within the Diapherotrites (detailed tree in **Fig. S5**).

Notably present in the genome is a nucleotide salvage pathway that features a form II/III RuBisCO (**Fig. S4**). The nucleotide salvage pathway generates G3P that can be converted to G2P, PEP, pyruvate, acetyl-CoA and acetate, with production of ATP. This is the only energy-generating step that could be identified in the genome. Ferredoxin and desulfoferredoxin are encoded in the genome but the mechanisms to reduce them are unknown. Reduced ferredoxin is required by pyruvate-ferredoxin oxidoreductase, a central component of the catabolic pathway and a key to metabolism, given the location of the genes involved near the origin of replication (Bryant et al. 2014; Chandler & Pritchard 1975). Encoded in the genome were also five large (>1800 aa) membrane-anchored proteins (**Fig. 3A**) within a single 42-kb region (5% of the entire genome) likely involved in cell-cell interaction. In fact, 10% of the proteome is encoded by enzymes of >1000 amino acids.

The 16S rRNA gene of the archaeon studied here places within the Diaphrerotrites, possibly representative of a new Class in this phylum (**Fig. 3B** and **Fig. S5**). We propose the name Candidatus “Forterrea multitransposorum” (after Patrick Forterre, for his research on archaea and archaeal replication). This novel archaeon appears to have a DNA-centric lifestyle and it may contribute to the critical task of carbon compound recycling in its nutrient-limited deep subsurface environment.

Previously described members of the phylum Diapherotrites supposedly acquired the genes that are necessary for a host-independent lifestyle from other organisms, likely bacteria, via horizontal gene transfer. Ca. “F. multitransposorum” is basal on the 16S rRNA gene tree to all these Diapherotrites and is predicted to have a host-dependent lifestyle. Given the fact that Ca. “F. multitransposorum” displays high genome fluidity via homologous recombination, it can be speculated that this mechanism is present in many Diapherotrites. Uptake of externally derived DNA may be responsible for the acquisition of bacterial genes enabling some of the cells to grow independent of their host.

## Data availability

The genome and the sequencing reads were deposited under NCBI BioProject number PRJNA405984 and the genome is available via http://ggkbase.berkeley.edu/Forterrea/organisms. The mapped read file can also be downloaded from this site under associated files.

## Acknowledgements

This study was funded by the Sloan Foundation (“Deep Life”, grant no. G-2016-20166041). AJP was supported by the German Research Foundation (DFG PR1603/01). Sequencing for this project was provided by the Emerging Technologies Opportunity Program of the U.S. Department of Energy Joint Genome Institute (Contract No. DE-AC02-05CH11231). We are grateful to scientists contributing to the sampling of the Crystal Geyser, particularly Karthik Anantharaman, Christopher Brown, Cathryn Ryan, and Bethany Ladd.

## Footnotes

Current address: University of Duisburg-Essen, Biofilm Centre, Group for Aquatic Microbial ecology, Universitaetsstrasse 5, 45141 Essen, Germany

